# Dosing Time Matters

**DOI:** 10.1101/570119

**Authors:** Marc D. Ruben, David F. Smith, Garret A. FitzGerald, John B. Hogenesch

## Abstract

Trainees in medicine are taught to diagnose and administer treatment as needed; time-of-day is rarely considered. Yet accumulating evidence shows that ∼half of human genes and physiologic functions follow daily rhythms. Circadian medicine aims to incorporate knowledge of these rhythms to enhance diagnosis and treatment. Interest in this approach goes back at least six decades, but the path to the clinic has been marked by starts, stops, and ambiguity. How do we move the field forward to impact clinical practice? To gain insight into successful strategies, we studied the results of more than 100 human trials that evaluated time-of-administration of drugs.

## Introduction

Trainees in medicine are taught to diagnose and administer treatment as needed; time-of-day is rarely considered. Yet a wealth of accumulating evidence shows that molecular clocks orchestrate 24 h rhythms in vital cardio-metabolic, endocrine, immunologic, and behavioral functions. Half of the mammalian protein-coding genome is expressed with a circadian rhythm (*1*, *2*). Time, therefore, adds a potentially important dimension to medicine.

Circadian medicine aims to incorporate knowledge of 24 h biological rhythms to enhance diagnosis and treatment. There are two broad strategies: 1) target the molecular clock, or 2) exploit its rhythmic outputs. In the former, the therapeutic target is the molecular oscillator itself. Light therapy for sleep disorders is the hallmark example, but feeding (*3*), exercise, and drugs (*4*, *5*) can also modulate the clock to potentially influence health (*4*). Our research focuses on the latter strategy – timing the delivery of treatment to coincide with rhythmic outputs of the clock. Interest in this approach goes back at least six decades (*6*), but the path to the clinic has been marked by starts, stops, and ambiguity. A series of recent insights, however, has renewed clinical interest.

Here, we discuss how current technology is shaping circadian medicine. Based on an analysis of more than 100 human studies, we provide recommendations for future trials in circadian medicine.

## Starts, stops, and scope

Fifty years ago, Shapiro and Rodwell showed that the rate-limiting enzyme in cholesterol synthesis oscillates with a 24 h period in rat liver (*7*, *8*). Fast-forward twenty years, and simvastatin — a short-acting inhibitor of this enzyme — was approved for the treatment of hyperlipidemia, labeled to take it “once a day in the evening”. This marked the first translation from knowledge of a circadian mechanism to a widely used medication.

In retrospect, simvastatin *might* have foreshadowed a wave of circadian drug development across therapeutic areas. It did not. Although studies in the mid-1990’s showed that rhythmically delivered chemotherapy could reduce toxicity in colorectal cancer (*9*–*11*), they did not lead to changes in FDA labeling, treatment guidelines, or standard of practice. Trials in a handful of other therapeutic areas around this time also supported time-dependent-dosing (*12*–*16*), with similarly limited influence. In fact, there are few examples of mechanism-based circadian medicine. Only 4 of the 50 currently most prescribed drugs (*17*) have FDA-labeled time-of-day dosing recommendation. The 20^th^ World Health Organization’s Model List of Essential Medicines makes no mention of dosing time.

### NEW DISCOVERIES RENEW CLINICAL INTEREST

Accumulating data show circadian regulation at genomic and physiologic levels that is far more extensive than previously understood. The central clock in the hypothalamus coordinates rhythmic behavior. However, local clocks exist in virtually every peripheral tissue in mammals (*18*). In a high-resolution time-series study of twelve mouse organs, nearly half of protein-coding genes were expressed with a 24 h rhythm in at least one organ (*1*). These clock-driven rhythms in gene expression generate rhythms in hormone secretion, lipid and glucose metabolism, lung and cardiovascular physiology, and many other tissue-specific functions (*19*). Remote sensors and wearable technology have revolutionized data collection from humans “in the wild.” Using multiple devices, a small feasibility study uncovered daily variation in 62% of physiologic and behavioral measures from 6 healthy individuals (*20*).

Expression of many disease phenotypes is also time-dependent. The incidence of heart attacks, and symptoms in asthma, allergic rhinitis, cancer, arthritis, depression and suicidal intent, peptic ulcers, and pain all show time-of-day variation (*21*). Thousands of rhythmically-expressed genes metabolize, transport, or are the targets of drugs (*1*, *22*). For drugs with rapid pharmacokinetics (PK), circadian time could influence efficacy and/or toxicity and thus, their therapeutic index (*23*).

### HARMONIZING TREATMENT WITH MOLECULAR RHYTHMS

Animal experiments are defining different ways to leverage circadian time for therapeutic benefit. Timing the administration of a drug to coincide with peak levels of its physiologic target was effective in models of atherosclerosis (*24*) and obesity (*25*), to name a few. Alternatively, drug delivery can be timed to coincide with *trough* levels of an *undesired* target, successfully executed in a mouse model of pancreatic cancer (*26*). Another approach is to harmonize the administration of a drug with rhythms in its absorption, distribution, metabolism, or excretion. Recent work identified a 24 h rhythm in blood-brain barrier (BBB) permeability that influences efficacy of an antiepileptic drug in a Drosophila model of seizure (*27*).

All of this has generated renewed enthusiasm for circadian time as a parameter in medicine; in 2018 alone, 15 reviews were published with “chronotherapy” in the title/abstract. But how do we move the field forward to influence clinical practice? To gain insight into successful strategies, we studied the results of 106 trials that evaluated time-of-administration of drugs.

## 50 years of clinical trials — an analysis

We identified human trials where time-of-day drug administration was the specific point of investigation (Fig. 1 and Table S1). Our criteria were published studies that directly compared at least two different time-of-day treatment schedules and measured efficacy or toxicity. Altogether, these studies comprised 70 distinct drugs or combinations or medical procedures across 15 different therapeutic areas.

**Fig 1.**
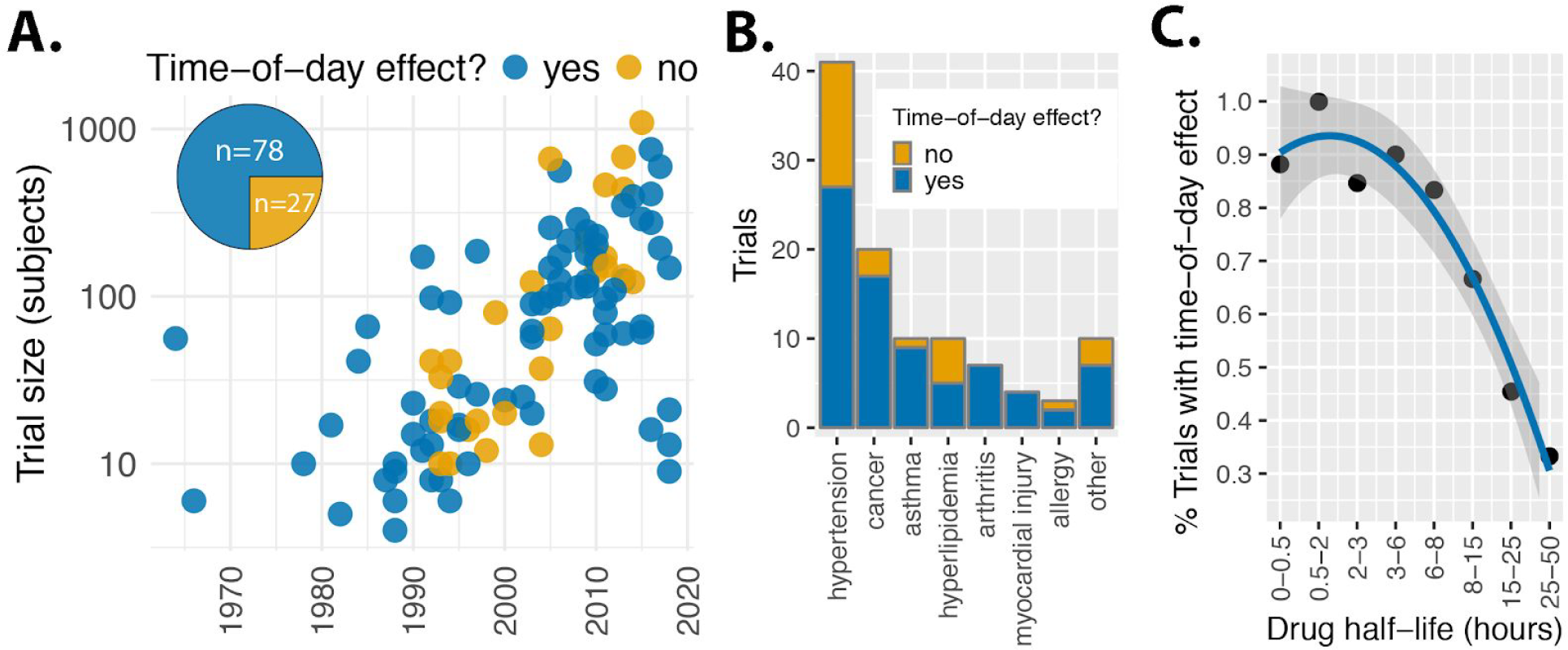
Circadian medicine is accelerating and has a track-record of success in highly prevalent diseases. **(A)** 75% of clinical trials (78/105) showed dosing-time-dependent efficacy or toxicity. **(B)** Numbers of trials by therapeutic area. Detailed information on drug type, half-life, and outcome for each trial shown in fig S1. **(C)** The relationship between drug half-life and time-of-day effect. Trials were grouped by drug half-life, with roughly equal numbers of trials in each group. For trials of combination therapies, we considered the drug ingredient with the shortest half-life.

### A TRACK-RECORD OF SUCCESS IN HIGHLY PREVALENT DISEASES

75% of trials (78/105) found that treatment efficacy or toxicity depended on dosing time (Fig 1A). Collectively, the work provides substantial support for circadian medicine in hypertension, cancer, asthma, and arthritis – diseases with extraordinarily high prevalence and unmet medical need (Fig 1B and fig S1). The number and size of trials in these areas has increased over the past several decades, reflecting well-known 24 h patterns in their phenotypic severity. Nevertheless, there are dozens of other indications (many with circadian components) and thousands of other approved drugs where the impact of time-of-day is unexplored.

### PHARMACOKINETICS MATTER, IN UNEXPECTED WAYS

Pharmacokinetics suggest that only drugs with half-lives of ~6 h or less would show time-of-day dependencies. Strikingly, our analysis shows that the majority of drugs with half-lives as long as 8–15 h were influenced by dosing-time (Fig 1C). This suggests that many commonly used drugs have narrow therapeutic windows and are influenced by dosing time.

The relationship described in Fig 1C helps to reconcile seemingly inconsistent trial results, even for drugs of the same class. For example, whereas *short*-acting simvastatin (half-life 3 h) was consistently more effective when taken in the evening (*28*–*31*), neither *long*-acting atorvastatin (*32*, *33*) (half-life 25 h) nor extended-release simvastatin (*34*, *35*) showed morning–evening differences in the majority of trials (fig S1). Long-acting formulations likely obscured circadian effects in other areas as well. Amolidipine, a long-acting Ca^2+^ channel blocker (half-life 40 h), did not lower blood pressure (BP) in a time-dependent manner (*36*, *37*). However, when it was combined with a short-acting angiotensin receptor blocker (valsartan or olmesartan) (*38*, *39*) or a diuretic (*40*), evening was more effective than morning dosing (fig S1).

Overall, 57 of 67 (85%) trials with half-lives < 15 h showed dosing time-dependence compared to just 9 of 23 (39%) for longer-acting drugs. This information could be the difference in an FDA approval.

### SMALL TRIALS AND TOO FEW TIME POINTS

There are still a number of discrepancies that cannot be explained by differences in drug kinetics (fig S1). Limitations in study design may contribute to this. Many of the trials were small. Median enrollment in the hypertension studies, for example, was 62 subjects, with only one study including more than 500 patients. By comparison, the most recently FDA-approved antihypertensive (Byvalson) was based on a pivotal trial of 4,161 hypertensives (*41*). This trial did not test for time-dependency. One of the components, valsartan, has a short half-life and established time-dependent efficacy (*42*–*44*).

Further, these studies were underpowered to detect periodic effects. The vast majority (~90%) compared only two time points, almost always morning versus evening, which might miss circadian effects at other times of day. Lastly, most studies did not account for possible interindividual variability in internal circadian phase, arising from differences in lifestyle, work schedule, or chronotype. 8AM on the wall clock is not “morning” for everyone.

Despite these limitations, there is significant support for circadian medicine. Yet, a major barrier to clinical translation has been the lack of understanding of how the circadian clock governs human physiology and pathophysiology. Even for medicines with demonstrable dosing-time-dependent effects, the mechanisms are not well defined.

## Present: towards mechanism-driven circadian medicine

### MAPPING DYNAMIC DRUG TARGETS IN HUMANS

Human time-series data have not been possible at the spatial and temporal resolution of animal models (*45*–*47*). Although longitudinal studies detected rhythmic gene expression from human samples (*48*–*52*), taking multiple biopsies from a patient over a 24 h period is difficult to execute, costly, and limited to the most easily accessible tissues. Alternatively, population-scale rhythms can be detected from single donor samples (*53*); however, time of sample collection is often not recorded. An algorithm called CYCLOPS was developed to reconstruct sample order in the absence of sample collection times (*26*).

CYCLOPS was used to build a first population-scale human atlas of circadian gene expression (2) and showed that regardless of a person’s sex, age, or ethnicity, the body’s internal clock regulates half of the protein-coding genome. This work connected thousands of different drugs – both investigational and approved – to thousands of cycling genes that encode drug targets, transporters, or metabolizing enzymes. This identified many specific mechanisms by which time-of-day might influence drug activity.

### TIME FOR STANDARD-OF-CARE THERAPIES

Genes that encode targets of antihypertensives cycle in the human cardiovascular system (*2*). These drugs are influenced by time of administration (Fig 1, fig S1). With knowledge of the *where* and *when* of drug target abundance, researchers and developers can make predictions about the influence of timed-dosing of many routinely prescribed short-acting medications.

Current knowledge might also help mitigate drug-related toxicity. Cardiotoxicity is the leading cause of drug discontinuation at all stages (*54*). Many drug classes, while not necessarily indicated for cardiovascular disease, actively target the heart and peripheral vasculature. For example, theophyllines –– bronchodilators administered for pulmonary disease –– function by inhibiting genes that are also critical to normal heart function (*55*–*57*). We now know that several of these “off-targets” are rhythmically expressed in human heart and vasculature (*2*). Can dosage time for these and other compounds be leveraged to reduce cardiotoxicity?

Time matters for several non-drug interventions as well. At least eight separate trials have shown that radiation therapy is less toxic when administered at a particular time (fig S1), thought to be when non-tumor cells are less susceptible to radiation damage (*58*). Circadian-dependent vulnerability to injury also exists in the context of cardiovascular procedures (*59*, *60*). Even with this understanding, practical limitations make implementation of timed-therapy difficult. Despite substantial support for timed administration of chemotherapy, this is not standard of practice due to multiple barriers (i.e. scheduling infusions at specific times and limited infrastructure).

## Future: drug development to drive circadian medicine

Ultimately, trials in circadian medicine will need to demonstrate an unambiguous health benefit over standard-of-care. Based on analysis of an ~50 year history of more than 100 clinical trials, we offer a set of recommendations for future studies in circadian medicine.

### MECHANISM FIRST

Genome-scale maps of 24 h rhythms are providing volumes of data for hypothesis-driven circadian medicine. There are ~2,000 approved drugs. Those whose targets oscillate in humans are a good place to start. Still, the best targets and times for treatment in the context of disease need to be determined empirically. The most effective medicines, we suspect, will incorporate knowledge of 24 h dynamics in both health and disease.

### SHORTER-ACTING AGENTS

Since the majority of support for circadian medicine comes from trials of drugs with half-lives < 15 h, we suggest that future efforts focus there. Although some very long-acting agents showed administration-time-dependent effects (fig S1, Table 1), the mechanisms are unclear. Trials should strive to delineate drug PK, ideally at more than one dosage time.

The industry trend towards sustained-release formulations, while good for patient adherence, can be problematic from the vantage of circadian biology. Drug exposure is constant over 24 h but its target(s) may not be. Further, flattening a process that is normally rhythmic may be maladaptive. For example, in people with high BP, short-acting antihypertensives were more effective than long-acting forms at restoring the normal nighttime drop in BP (*61*) – which may improve cardiovascular outcomes (*62*).

### BEYOND MORNING VERSUS EVENING

The simplest comparison, between morning and evening dosing (91 of 105 trials), can only confirm or rule out a difference between these two times. Although medications are typically taken at these times for convenience, other times may be more effective or safe. For drugs with short half-lives, two-time point sampling is likely insufficient. While the gold standard of hourly sampling for genome-scale circadian analysis (*63*) may be impractical for human trials, clinical researchers should nevertheless weigh the upstream cost of additional time-points against the downstream cost of a type II error.

### BIOMARKING INTERNAL TIME

Trials in circadian medicine must control for interpersonal differences in phase. At a minimum, subject’s rest-activity rhythms provide a reasonable estimate of internal circadian phase.

Recently, three independent studies reported algorithms capable of inferring circadian phase from skin (*64*) or blood samples (*65*, *66*). These pave the way for development of biomarkers to help optimize the delivery time of treatment for patients where determining circadian phase may be difficult (e.g. patients in an ICU).

### CONTROL FOR CIRCADIAN DISRUPTION

Many of the trials we studied were conducted in artificial environments. Subjects in inpatient care settings — often detached from natural environmental fluctuations for extended periods of time — are at risk for circadian disruption. Simply recreating changes in natural light over 24 hours improved recovery in experimental (*67*) and hospital-based clinical studies (*68*–*70*). Besides light, noise levels and frequent awakenings are also sources of circadian disruption.

This becomes a critical consideration for time-of-day effects and is leading many to rethink hospital layouts from the vantage of circadian biology.

## Conclusions

The most commonly prescribed drugs help only a minority of patients who take them (*71*). Although the common refrain to this is a call for better ***genetic*** **precision**, circadian effect sizes can be just as large. **Circadian precision** medicine aims to deliver treatment in harmony with target physiology and has shown that it can improve efficacy and safety in many feasibility-sized trials in hypertension, arthritis, asthma, and cancer, among others. Recent technological advances in circadian biology have the potential to exert broad influence on medical practice.

At the time of this review, there is ***one*** registered clinical trial in the United States with time-of-day endpoints: *Temozolomide Chronotherapy for High Grade Glioma* (*72*). Will circadian biology factor into the future of treatment? Time will tell.

---

**Fig S1.**
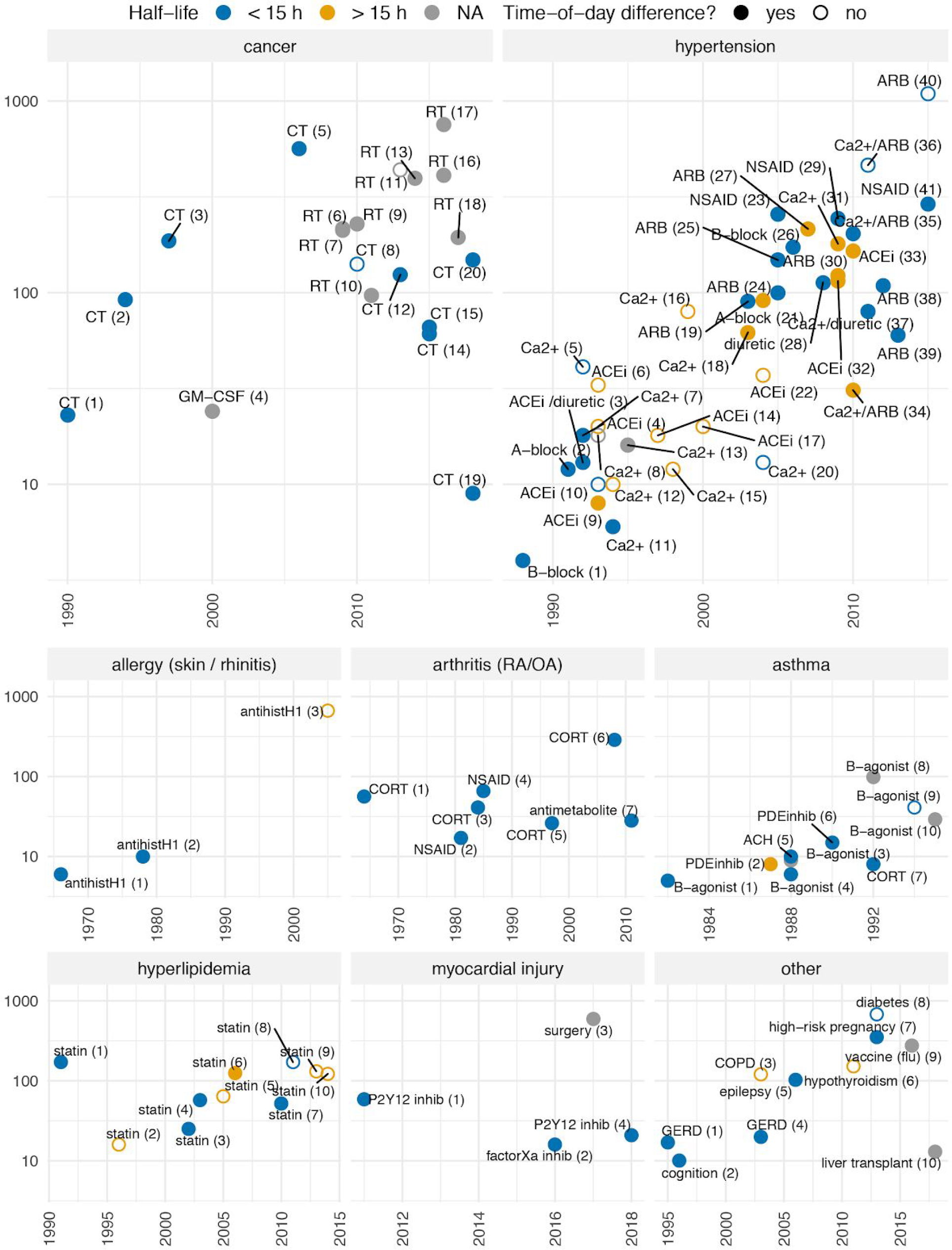
50 years of clinical studies testing dosing-time-dependence of medical treatment. 105 clinical studies that were: 1) published, 2) directly compared at least two different time-of-day treatment schedules, and 3) measured clinical effect or toxicity. Point color indicates drug half-life: short (≤15 h), long (>15 h), or unknown (NA). Point fill distinguishes trials that found significant (P-value < 0.05) administration-time-dependent efficacy and/or toxicity (solid) from trials that did not (empty). Trials grouped by therapeutic area and labeled according to drug class. Study citations are indicated by the number in parentheses for each trial (table S1).

**Table S1.**
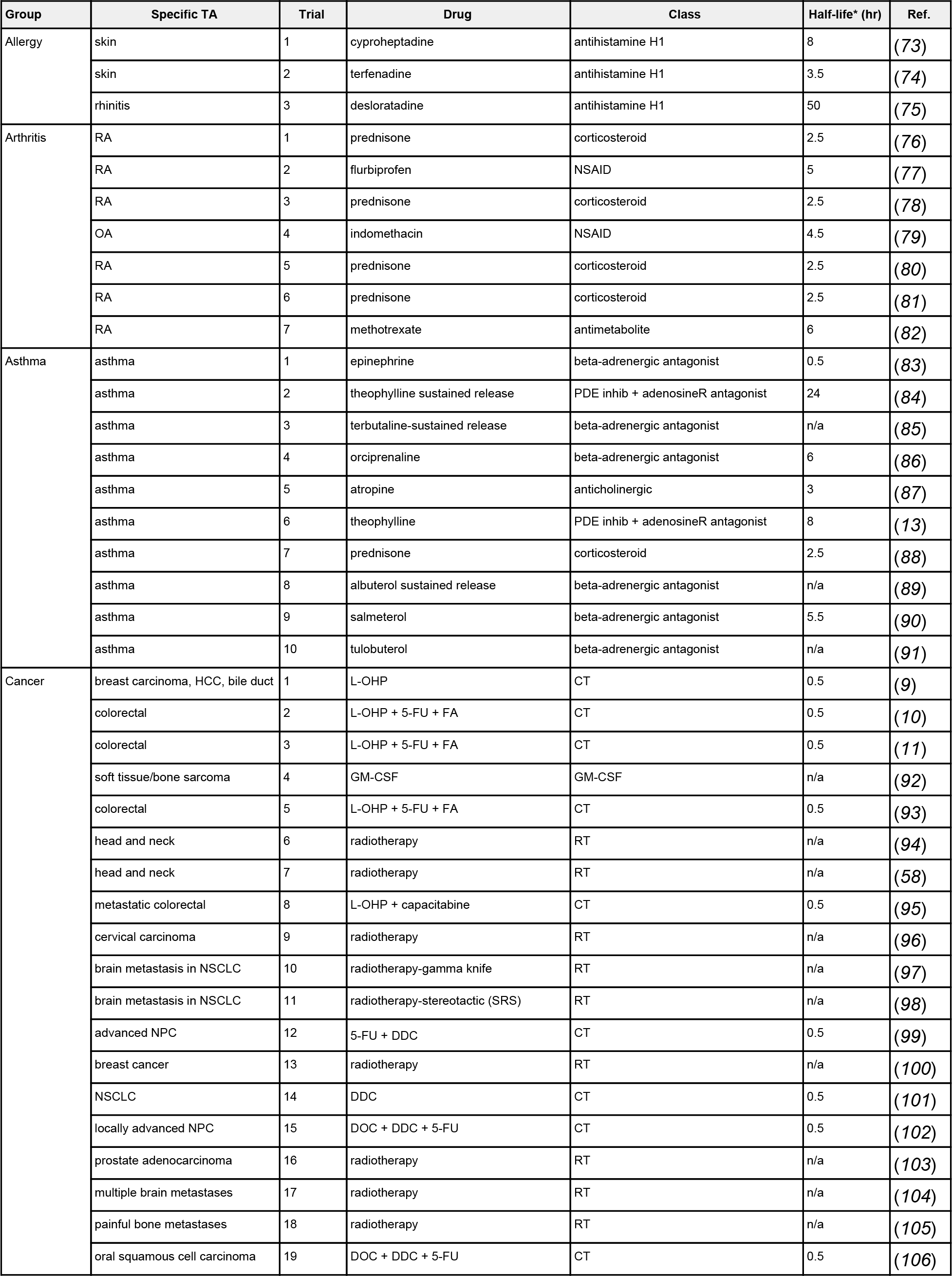
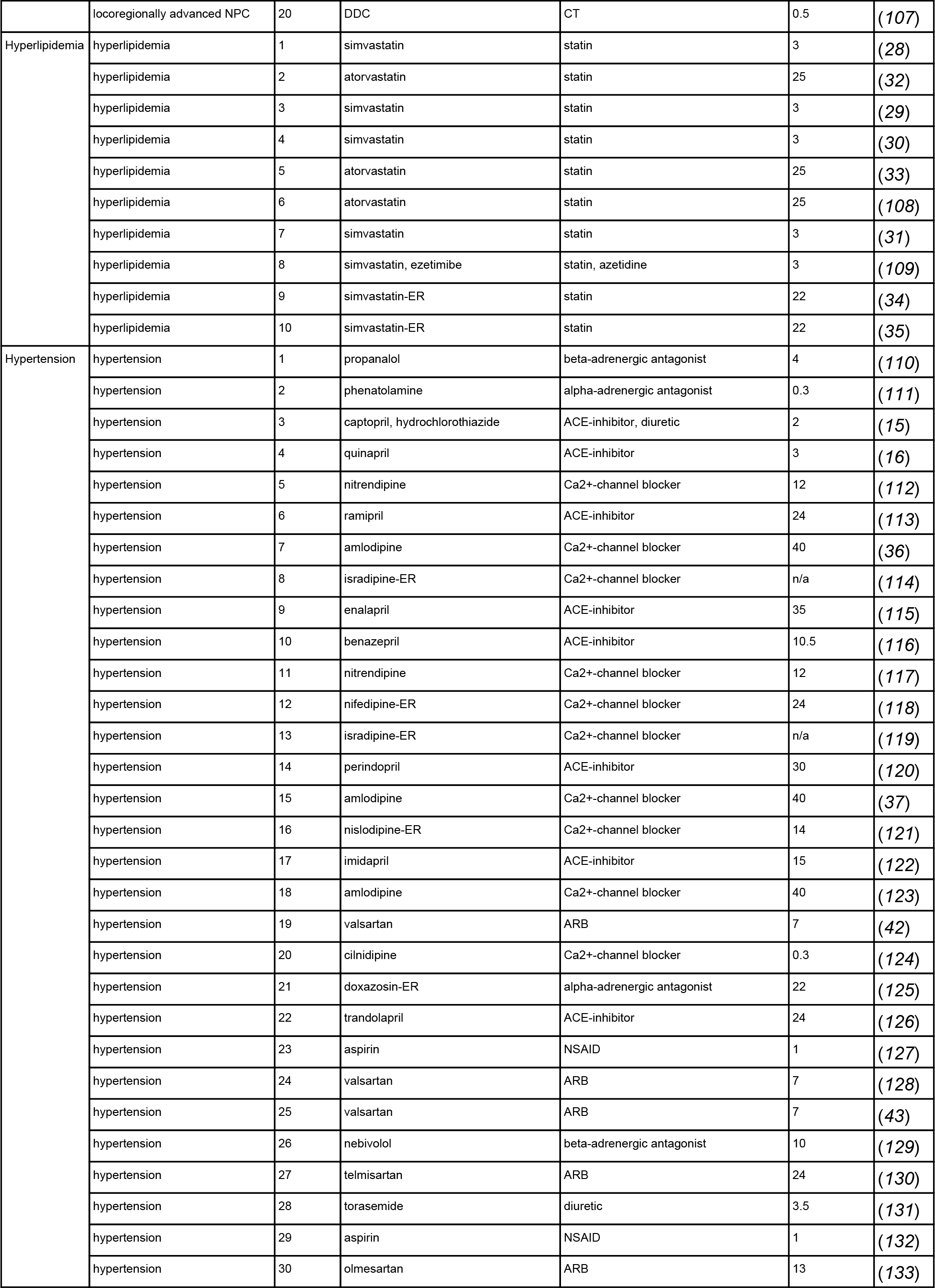
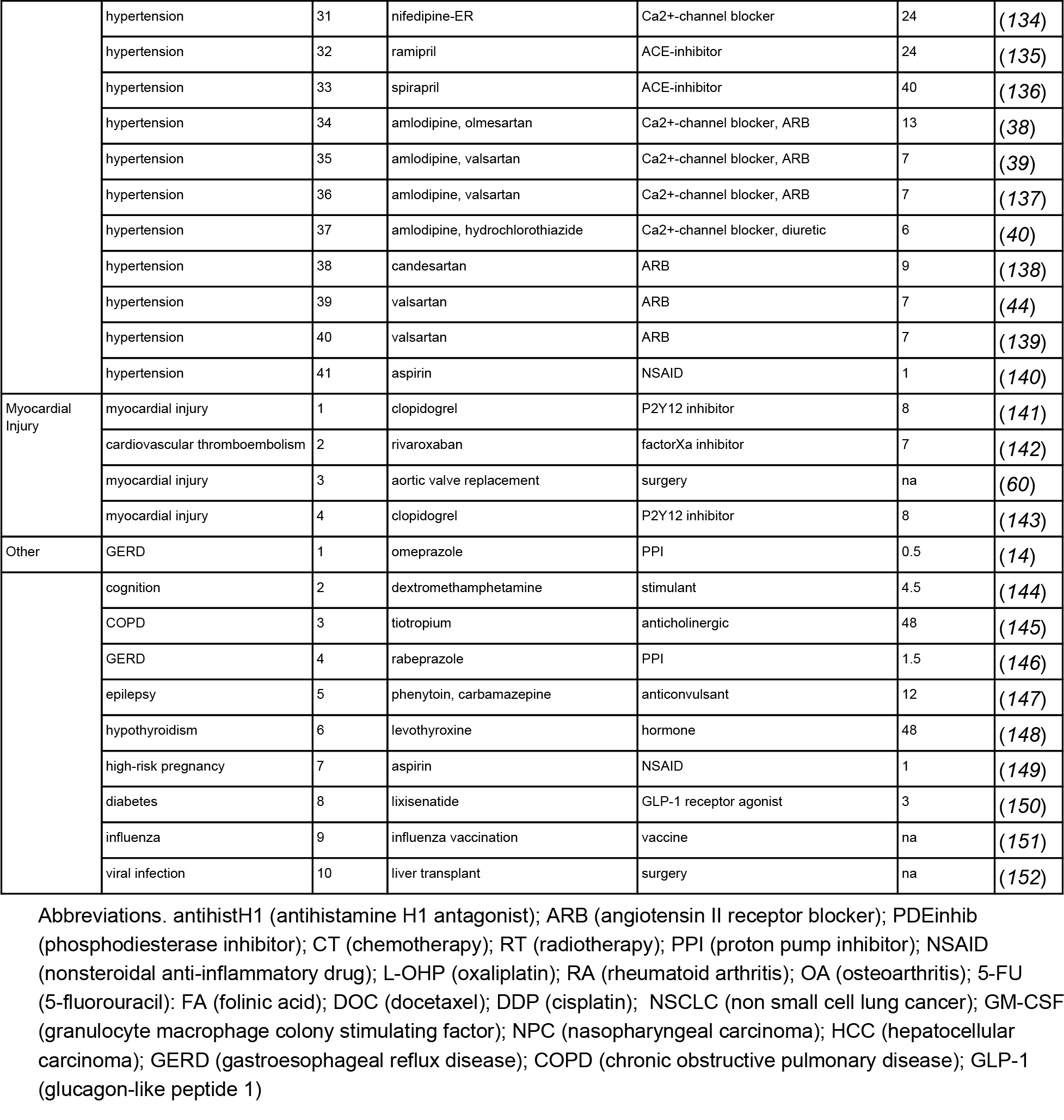
Details for each trial and reference. *Half-lives from DrugBank 5.0 (*153*), FDA-label information, or research publications. For drug combinations, the shortest-acting agent in the combination is reported in this table and in Fig 1.

## Author contributions

M.D.R. and J.B.H. designed research; M.D.R. and D.F.S. performed research; M.D.R, D.F.S., G.A.F., and J.B.H. wrote the paper.

